# Effect of chronic JUUL aerosol inhalation on inflammatory states of the brain, lung, heart and colon in mice

**DOI:** 10.1101/2021.03.09.434442

**Authors:** Alex Moshensky, Cameron Brand, Hasan Alhaddad, John Shin, Jorge A. Masso-Silva, Ira Advani, Deepti Gunge, Aditi Sharma, Sagar Mehta, Arya Jahan, Sedtavut Nilaad, Daniyah Almarghalani, Josephine Pham, Samantha Perera, Kenneth Park, Rita Al-Kolla, Hoyoung Moon, Soumita Das, Min Byun, Zahoor Shah, Youssef Sari, Joan Heller Brown, Laura E. Crotty Alexander

## Abstract

While health effects of conventional tobacco are well defined, data on vaping devices, including the most popular e-cigarette JUUL, are less established. Prior acute e-cigarette studies demonstrated inflammatory and cardiopulmonary physiology changes while chronic studies demonstrated extra-pulmonary effects, including neurotransmitter alterations in reward pathways. In this study we investigated effects of chronic flavored JUUL aerosol inhalation on inflammatory markers in brain, lung, heart, and colon. JUUL induced upregulation of cytokine and chemokine gene expression and increased HMGB1 and RAGE in the nucleus accumbens. Inflammatory gene expression increased in colon, and cardiopulmonary inflammatory responses to acute lung injury with lipopolysaccharide were exacerbated in the heart. Flavor-dependent changes in several responses were also observed. Our findings raise concerns regarding long-term risks of e-cigarette use as neuroinflammation may contribute to behavioral changes and mood disorders, while gut inflammation has been tied to poor systemic health and cardiac inflammation to development of heart disease.

**One Sentence Summary:** Chronic, daily inhalation of pod-based e-cigarette aerosols alters the inflammatory state across multiple organ systems in mice.

## Introduction

Chronic inhalation of tobacco smoke is known to damage multiple cell types and cause a wide range of diseases throughout the body. In particular, many pulmonary inflammatory diseases are caused and affected by conventional tobacco use (1, 2). It is also known that nicotine affects brain development and alters responses to addictive substances. Nicotine activates carcinogenic pathways, putting users at an increased risk of cancer (3). With unproven claims to be a safer alternative to cigarette smoking, modern electronic (e)-cigarette devices were introduced in 2003 as a novel nicotine delivery system (4, 5). The JUUL^TM^, a device that gained popularity due to its sleek and concealable design, has utilized pods containing e-liquids with enticing flavors such as Mango, Mint, and Crème Brulee (now discontinued)(6). However, the health effects of chronic inhalation of aerosols generated from pod devices remain largely unknown.

While the data on health effects of conventional tobacco are extensive, the data on e-cigarettes and vaping devices are less established due to their recent entry to the market (7, 8). In particular, research in this area is impeded by the rapid evolution of vaping devices. The vape pens and cig-a-likes were the first e-cigarettes studied from 2007-2014, whereas the box Mods became highly popular and research on these devices began around 2015. Pod devices, including the JUUL, were invented in 2016 and rapidly dominated the market by 2017-2020, however, studies on the harmful effects of these types of vaping devices are scarce (9). Thus, research on the health effects of these pod-based devices, which produce aerosols with a different chemical composition than prior devices (including often significant higher concentrations of nicotine than Mod devices), is desperately needed.

Because of the short time e-cigarettes and vaping devices have been on the market, very little is known about the long-term chronic effects of vaping. Acute and subacute studies in human subjects have demonstrated changes in lung and cardiac function, with increased airway reactivity and lung inflammation, and increased heart rate and blood pressure in response to vaping (7). Previous studies of chronic effects of vaping are limited to e-cigarette aerosol inhalation models in animals but have demonstrated more profound effects, including renal, cardiac and liver fibrosis (10), emphysema (11), lung cancer (12), increased lung injury in the setting of influenza infection (13), increased arterial stiffness and atherosclerosis, and activation of addiction neurocircuits in the brain (14, 15).

The health effects of vaping JUUL pods remain unknown, despite the popularity of pod-based e-devices. Here, we broadly assessed the effects of daily JUUL aerosol inhalation on cardiopulmonary function and inflammation across organ systems, including the reward pathways in the brain. We induced acute lung injury with inhaled *E. coli* lipopolysaccharide (LPS) to determine whether chronic JUUL use predisposes to deleterious responses in the setting of common infectious challenges such as Gram-negative bacterial pneumonia. We demonstrated here that daily inhalation of JUUL aerosols can lead to inflammatory changes in the brain, heart, lung, and colon, as well as alterations in physiological functions.

## Results

### Chronic JUUL inhalation associated with neuroinflammation in the brain

Previous studies have shown that conventional tobacco smoking increases proinflammatory cytokines in the brain, specifically TNFα, IL-1β, IL-6 (16, 17). Therefore, gene expression of these inflammatory cytokines were measured by qPCR in different brain regions of mice exposed to JUUL Mango and JUUL Mint, as well as Air controls for 1 or 3 months. Specifically, we assessed gene expression in the nucleus accumbens core and shell (NAc-core and NAc-shell), and hippocampus, regions involved in behavior modification, formation of drug reward and anxious or depressive behaviors, and learning and memory, respectively. We observed that TNFα gene expression was significantly increased in the NAc-core and NAc-shell of mice exposed to 1 or 3 months of JUUL Mango or JUUL Mint compared to Air controls (**Figure 1A–D**). In contrast, TNFα levels were unchanged in the hippocampus throughout the exposures (**Figure 1E–F).** IL-1β gene expression was also significantly elevated in JUUL Mint- and Mango-exposed mice group in both the NAc-core and NAc-shell at 1 month compared to air control group (Figure 1G and 1I), but remained elevated at 3 months only in the NAc-shell (Figure 1J) and not in NAc-core (Figure 1H). The hippocampus showed unchanged levels of IL-1β gene expression at 1 and 3 months across groups (Figure 1K and 1L). In the case of IL-6, we observed a significant increase in gene expression in the NAc-shell in both JUUL Mango and JUUL Mint groups at 1 and 3 months (**Figure 1O and 1P**), but no significant differences were observed in the NAc-core and hippocampus when compared to Air controls (**Figure 1M, 1N, 1Q and 1R**). Overall, these data suggest that exposure to JUUL Mint and JUUL Mango may induce neuroinflammation in brain regions responsible for behavior modification, drug reward and formation of anxious or depressive behaviors (18).

**Figure 1.**
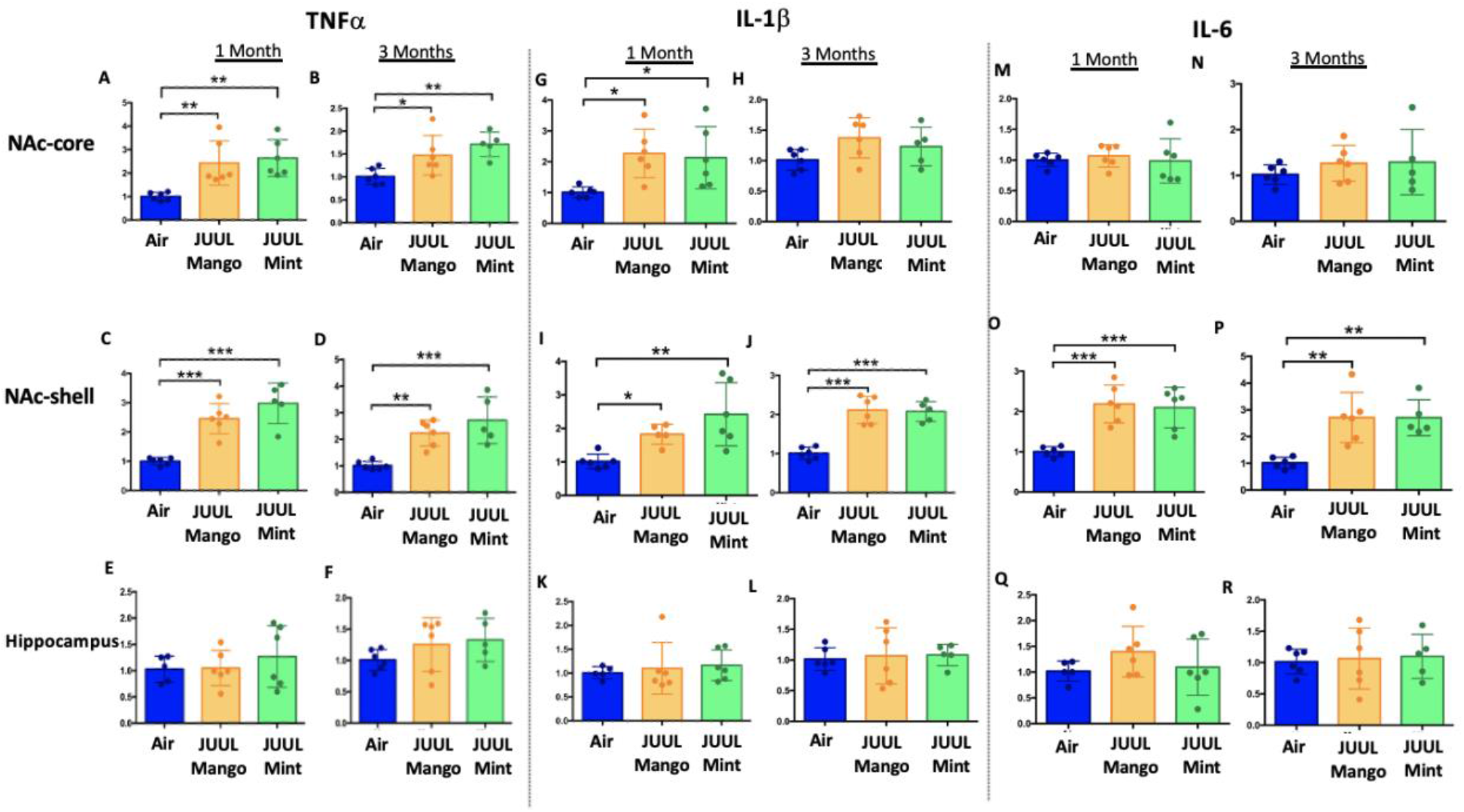
Chronic JUUL use leads to an increase of pro-inflammatory cytokines in different regions of the brain. Brains were harvested at the end point and the regions for NAc-core, NAc-shell and Hippocampus were sectioned. Later, the RNA was extracted and qPCR was performed to quantify the expression of TNFα, IL-1β, IL-6. TNFα expression is shown from NAc-core at A) 1 month and B) 3 months, from NAc-shell at C) 1 month and D) 3 months, and from Hippocampus at E) 1 month and F) 3 months. IL-1β expression is shown from NAc-core at G) 1 month and H) 3 months, from NAc-shell at I) 1 month and J) 3 months, and from Hippocampus at K) 1 month and L) 3 months. IL-6 expression is shown from NAc-core at M) 1 month and N) 3 months, from NAc-shell at O) 1 month and P) 3 months, and from Hippocampus at Q) 1 month and R) 3 months. Data are presented as individual data points ± SEM with n=5-6 mice per group. \**p*<0.05, \*\**p*<0.01, *** *p*<0.001 and **** *p*<0.0001.

To further confirm the neuroinflammatory response associated with chronic JUUL exposure, we measured levels of receptors for advanced glycation end products (RAGE) and its ligand high mobility group box 1 (HMGB1) protein by western blot in the NAc-core, NAc-shell, and hippocampus of mice exposed to JUUL Mango, JUUL Mint and Air at 1 and 3 months. RAGE and HMGB1 have been implicated in inducing neuroinflammation (16), and previous studies have shown that HMBG1-1 and RAGE expression are increased with exposure to cigarette smoke (19, 20). No significant changes of HMGB1 were observed in NAc-core at 1 or 3 months of JUUL aerosol exposure between groups (**Figure 2A-B)**. The NAc-shell, however, showed significant increase in HMGB1 at 1 and 3 months in mice exposed to JUUL Mango and JUUL Mint relative to Air controls (**Figure 2C–D),** and the increase was more pronounced at 3 months (**Figure 2D**). The hippocampus showed no changes in HMGB1 protein expression at 1 month (**Figure 2E**), and actually showed significant decrease in protein expression in mice exposed for 3 months to either JUUL Mango or JUUL Mint as compared to Air controls (**Figure 2F**). In the case of RAGE, the protein levels were not significantly altered in all tested brain regions (**Figure 2G-I, 2K-L**), except in the NAc-shell of the mice exposed to JUUL Mango or JUUL Mint for 3 months when compared to Air controls (**Figure 2J**). Altogether, these data indicate that exposure to aerosols from JUUL devices induced neuroinflammation in reward brain regions, particularly that of the NAc-shell and NAc-core regions.

**Figure 2.**
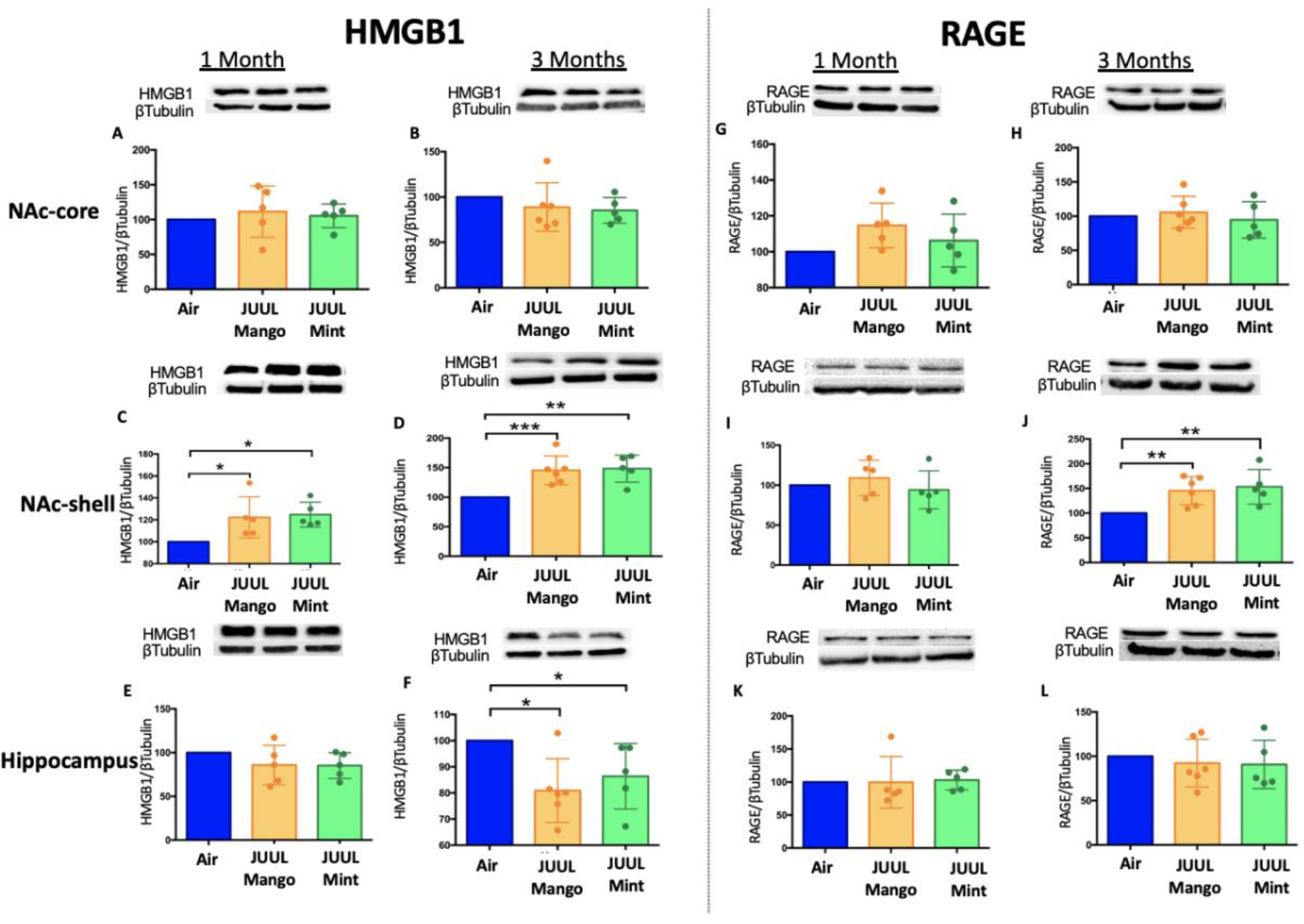
Chronic JUUL use leads to an increase of inflammatory mediators HMGB1 and RAGE. Brains were harvested at the end point and the regions for NAc-core, NAc-shell and Hippocampus were sectioned. Later, protein was extracted and Western Blot was performed to quantify the expression of HMGB1-1 and RAGE. HMGB1-1 relative protein level are shown from NAc-core at A) 1 month and B) 3 months, from NAc-shell at C) 1 month and D) 3 months, and from Hippocampus at E) 1 month and F) 3 months. RAGE protein levels are shown from NAc-core at G) 1 month and H) 3 months, from NAc-shell at I) 1 month and J) 3 months, and from Hippocampus at K) 1 month and L) 3 months. Changes in proteins levels are relative to Air controls. Data are presented as individual data points ± SEM with n=5-6 mice per group. \**p*<0.05, \*\**p*<0.01 and *** *p*<0.001.

### Chronic inhalation of JUUL aerosols alters inflammatory and fibrosis associated gene expression in cardiac tissue

Changes in the myocardium have been widely observed in response to cigarette smoking, and we previously showed that inhalation of e-cigarette aerosol generated by Vape pens for 3-6 months induced fibrotic changes in cardiac tissue (10). Fibrosis is typically driven by either cellular injury or inflammation. Increases in pro-inflammatory cytokines and fibrosis-associated proteins have been linked to the development of cardiovascular diseases (21–24). Based on the inflammatory and fibrotic markers commonly observed in response to myocardial infarction and development of heart failure, we assessed the expression of mRNA for TNFα, IL-1β IL-6, IL-18, CCL2, CCL3, CXCL1, CXCL2, Col1a1, Col3a1, Postn and TLR4 at 1 and 3 months **(Figure 3A-X**). Surprisingly, none of the pro-inflammatory cytokines or chemokines examined were upregulated by JUUL exposure. Indeed, TNFα, IL-6, and CXCL2 were actually down-regulated in 1 month JUUL Mint exposed mice, as was the pro-fibrotic gene Col1a1 (**Figure 3A, 3E, 3O and 3Q**). In contrast to JUUL Mint, aerosol inhalation of JUUL Mango for 1 month was only associated with the downregulation of CXCL2 (**Figure 3O**). JUUL Mint and Mango aerosol inhalation also had differing effects on CXCL2 and TLR4 expression at 3 months of exposure (**Figure 3P and 3X, respectively**). These findings suggest that there may be flavor-specific effects in addition to nicotine-specific effects on the cardiovascular system. Overall, these changes show that inflammatory pathways in cardiac tissue are affected by JUUL aerosol inhalation. While overt inflammation is not induced, it is well known that any alterations to the immune-inflammation axis, activating or suppressive, can lead to changes in disease susceptibility and incidence (25).

**Figure 3.**
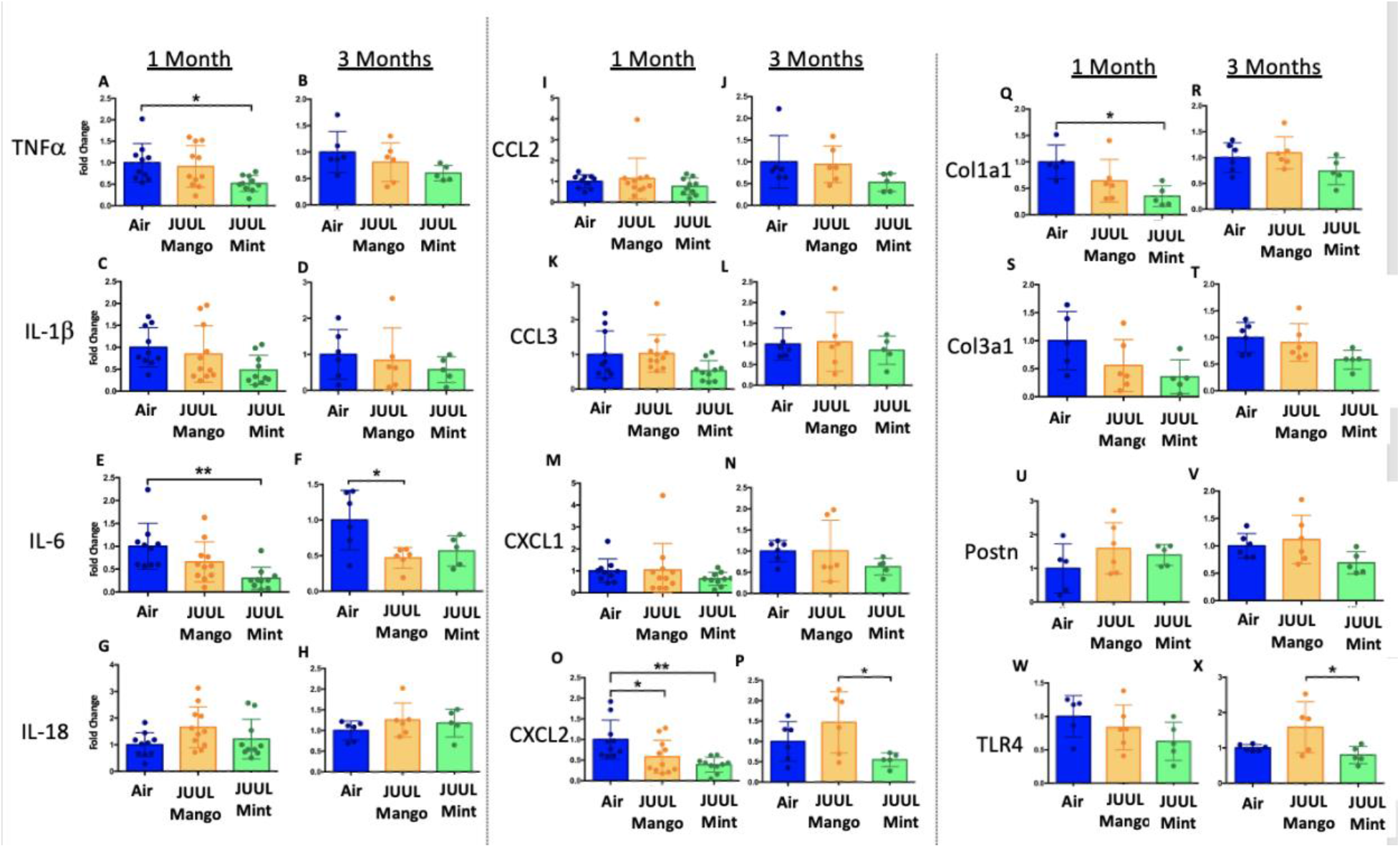
Chronic inhalation of JUUL aerosols alters inflammatory and fibrosis associated gene expression in cardiac tissue. Hearts were harvested, and RNA was extracted from the left ventricle and qPCR was performed to quantify the gene expression of different cytokines, chemokines and fibrosis-associated genes. Cytokines include TNFα at A) 1 month and B) 3 months, IL-1β at C) 1 month and D) 3 months, IL-6 at E) 1 month and F) 3 months, and IL-18 at G) 1 month and H) 3 months. Chemokines include CCL2 at I) 1 month and J) 3 months, CCL3 at K) 1 month and L) 3 months, CXCL1 at M) 1 month and N) 3 months, and CXCL2 at O) 1 month and P) 3 months. Fibrosis-associated genes include Col1a1 at Q) 1 month and R) 3 months, Col3a1 at S) 1 month and T) 3 months, Postn at U) 1 month and V) 3 months, and TLR4 at W) 1 month and X) 3 months. Changes in expression levels are relative to Air controls. Data are presented as individual data points ± SEM with n=5-11 mice per group. \**p*<0.05 and \*\**p*<0.01.

### Chronic JUUL aerosol inhalation alters pro-inflammatory markers in the colon

Due to the altered inflammation observed in brain and cardiac tissue, we also assessed inflammation in the colon, where cigarette smoking has been shown to alter inflammation and thereby promote chronic digestive diseases (26, 27). In terms of documenting the effects of e-cigarettes on gastrointestinal inflammation, our knowledge is limited to only the study done by our research group, with a focus on changes induced by aerosols generated by box Mods only (28). In order to assess JUUL induced changes in the gastrointestinal tract, we examined inflammatory gene expression at 1 and 3 months of JUUL exposure. JUUL Mango induced upregulation of TNFα, IL-6, and IL-8 relative to Air controls after 1 month exposure (**Figure 4A, 4C, 4G**). Interestingly, at 3 months, JUUL Mango treatment resulted in less expression of TNFα, IL-6 and IL-1β than that observed in Air controls or JUUL Mint exposed mice (**Figure 4B, 4D, 4F**), but increased expression of CCL2 (**Figure 4J**). These data suggest that exposure to JUUL Mango aerosols modulates inflammation in the colon, primarily inducing key inflammatory cytokines at 1 month. In JUUL Mango and Mint, there was no change in IL-1B or CCL2 at 1 month (**Figure 4E and 4I**) and IL-8 at 3 months (**Figure 4H**).

**Figure 4.**
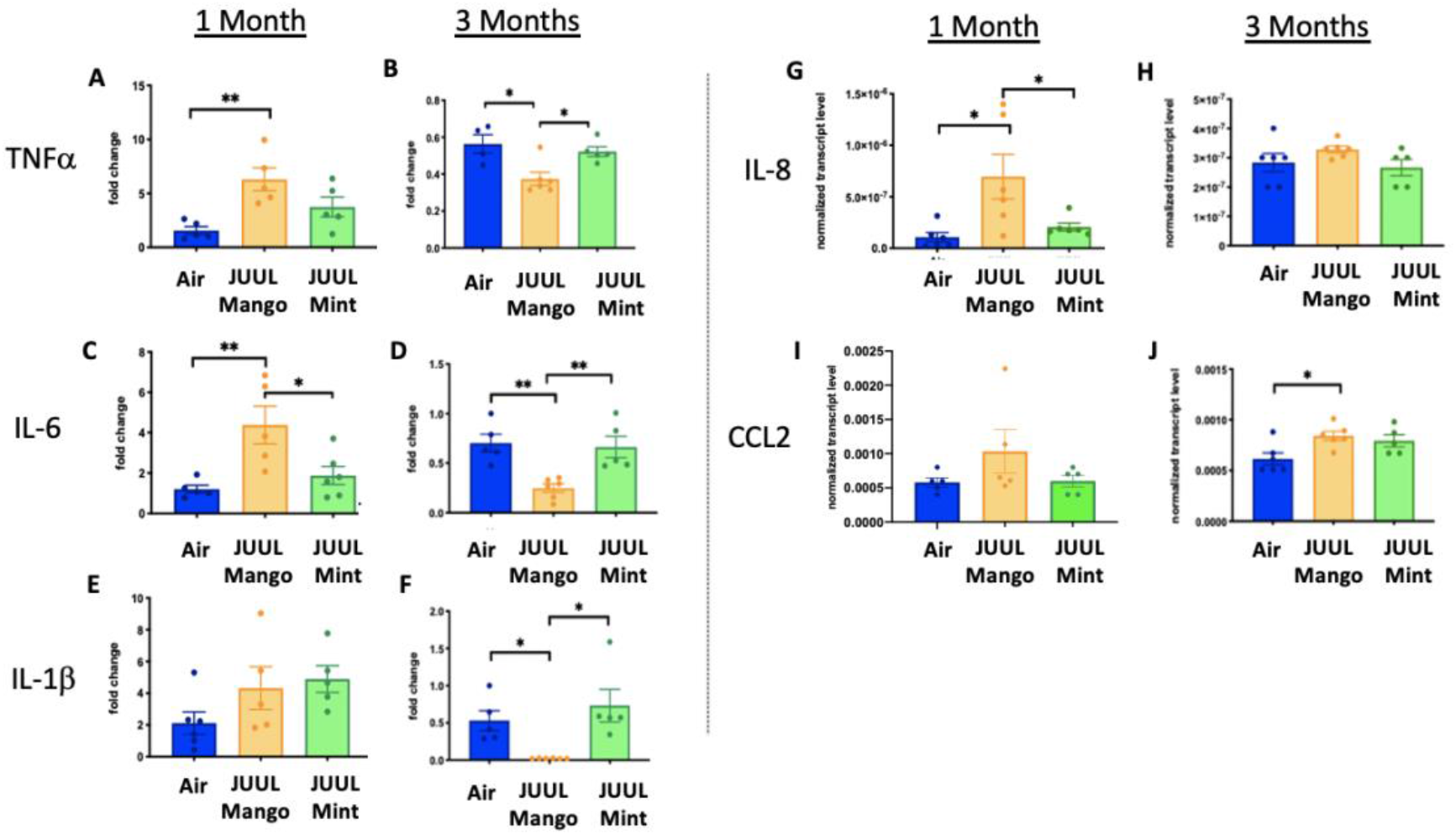
Chronic JUUL aerosol inhalation alters pro-inflammatory markers in colon. Inflammation was assessed in the colon at 1 and 3 months. Panels show inflammation markers in the colon in TNFα A) 1 month and B) 3 months, IL-6 at C) 1 month and D) 3 months, IL-1β at E) 1 month and F) 3 months, IL-8 G) and H), and CCL2 I) 1 month and J) 3 months. Data for inflammation markers is presented as individual data points ± SEM. * *p*<0.01 and ** *p*<0.001.

### Daily JUUL aerosol inhalation does not alter heart rate and blood pressure

Chronic exposure to cigarette smoke leads to cardiovascular changes, mediated through altered autonomic tone, but little is known about the chronic cardiovascular effects of e-cigarettes, especially with pod devices (7). Thus, we exposed mice to JUUL aerosols and carried out assessments of cardiovascular function, including blood pressure (BP), heart rate (HR), and heart rate variability (HRV). Heart rate variability was determined from root-mean square differences of successive R-R intervals (RMSSD) and the mean of the standard deviations for all R-R intervals (SDNN). There were no significant changes in HR or HRV at 1 and 3 months of either JUUL Mint or JUUL Mango exposure relative to Air controls (**Supplemental Figure 1**). Similarly, systolic and diastolic BP were also unchanged relative to Air controls at either 1 or 3 months (**Supplemental Figure 1**). Thus, chronic exposure to pod-based e-cigarette aerosols containing high levels of nicotine does not appear to alter normal physiological autonomic cardiovascular regulation in mice.

### Chronic inhalation of JUUL aerosols does not alter pulmonary physiology

Lungs represent the main site for aerosol deposition during inhalant use. Several studies have shown the effects of e-cigarettes on lung physiology (7, 29). To determine the effect of chronic JUUL aerosol inhalation on lung physiology, mechanic scans of airways resistance and lung elastance were carried out and were found to be similar across JUUL Mint, JUUL Mango and Air control groups at 1 and 3 months of exposure (**Supplemental Figure 2**). Pressure-volume (PV) loops also demonstrated similarities amongst the three groups at 1 and 3 months (**Supplemental Figure 2**). Airway hyperreactivity was tested by methacholine challenge and revealed no differences amongst groups, as measured at 1 and 3 months (**Supplemental Figure 2**). Thus, long-term exposure to JUUL Mint and Mango aerosols does not appear to lead to significant changes in airway physiology.

### Chronic JUUL inhalation does not change the inflammatory state of lungs under homeostatic conditions

Conventional tobacco as well as some e-cigarette aerosol exposures, have been found to cause increased cellularity in the airways (2, 7, 8), and cigarette smoking leads to recruitment of neutrophils to the airways in particular (2). Total leukocyte and neutrophil counts in bronchoalveolar lavage (BAL) of mice exposed to JUUL aerosols for 1 and 3 months were not different than those in Air controls, indicating that inflammatory cell recruitment to the airways was unaffected (**Supplemental Figure 3**). Moreover, fixed lung sections stained with H&E showed no difference in inflammation in the lungs at 1 and 3 months of JUUL aerosol exposure as compared to Air controls (**Supplemental Figure 3**).

### Chronic JUUL inhalation does not induce cardiac, renal or liver fibrosis

Our previous studies with mice exposed to aerosols generated from Vape pens not only found fibrosis in cardiac tissue after 3-6 months of e-cigarette aerosol inhalation but also in the liver and kidneys (10). Cigarette smoking is also known to cause organ fibrosis (30). There were, however, no significant changes in fibrosis assessed by quantification of collagen fibers stained with Masson’s trichrome, in the liver, heart or kidneys of mice that inhaled JUUL Mango or JUUL Mint aerosols for 3 months relative to Air controls (**Supplemental Figure 4**).

### Impact of JUUL exposure on airway inflammation in the setting of inhaled LPS challenge

Long-term cigarette smoking is known to predispose to greater inflammatory responses to lung infections (2). However, few studies have examined the effects of e-cigarette vaping on the severity of attendant respiratory diseases (13). Under homeostatic conditions, the BAL of JUUL Mango and JUUL Mint mice contains similar levels of inflammatory cytokines and chemokines at both 1 and 3 months (**Figure 5A-N**). Inhaled LPS is a model of Gram-negative bacterial pneumonia and acute lung injury in mice. Mice exposed to JUUL aerosols and challenged with inhaled LPS had similar total numbers of leukocytes and neutrophils in the airways relative to Air controls (**Supplemental Figure 3**), and histological analysis of H&E staining showed that parenchymal inflammation was similar across groups after LPS challenge (**Supplemental Figure 3**). LPS challenge also leads to increased levels of CCL2 and KC (murine homolog of IL-8) in the airways. The increases in CCL2 and KC elicited by LPS were diminished in mice exposed to JUUL Mint, demonstrating an attenuated inflammatory response to LPS after sub-acute exposure to JUUL (**Figure 5A and 5C**, respectively). Differences in LPS induced cytokine levels were no longer observed after 3 month JUUL exposure versus Air control groups (**Figure 5B, 5D, 5F, 5H, 5J, 5L and 5N**), suggesting that chronic use of JUUL does not significantly alter inflammatory responses to Gram-negative infections in the lung.

**Figure 5.**
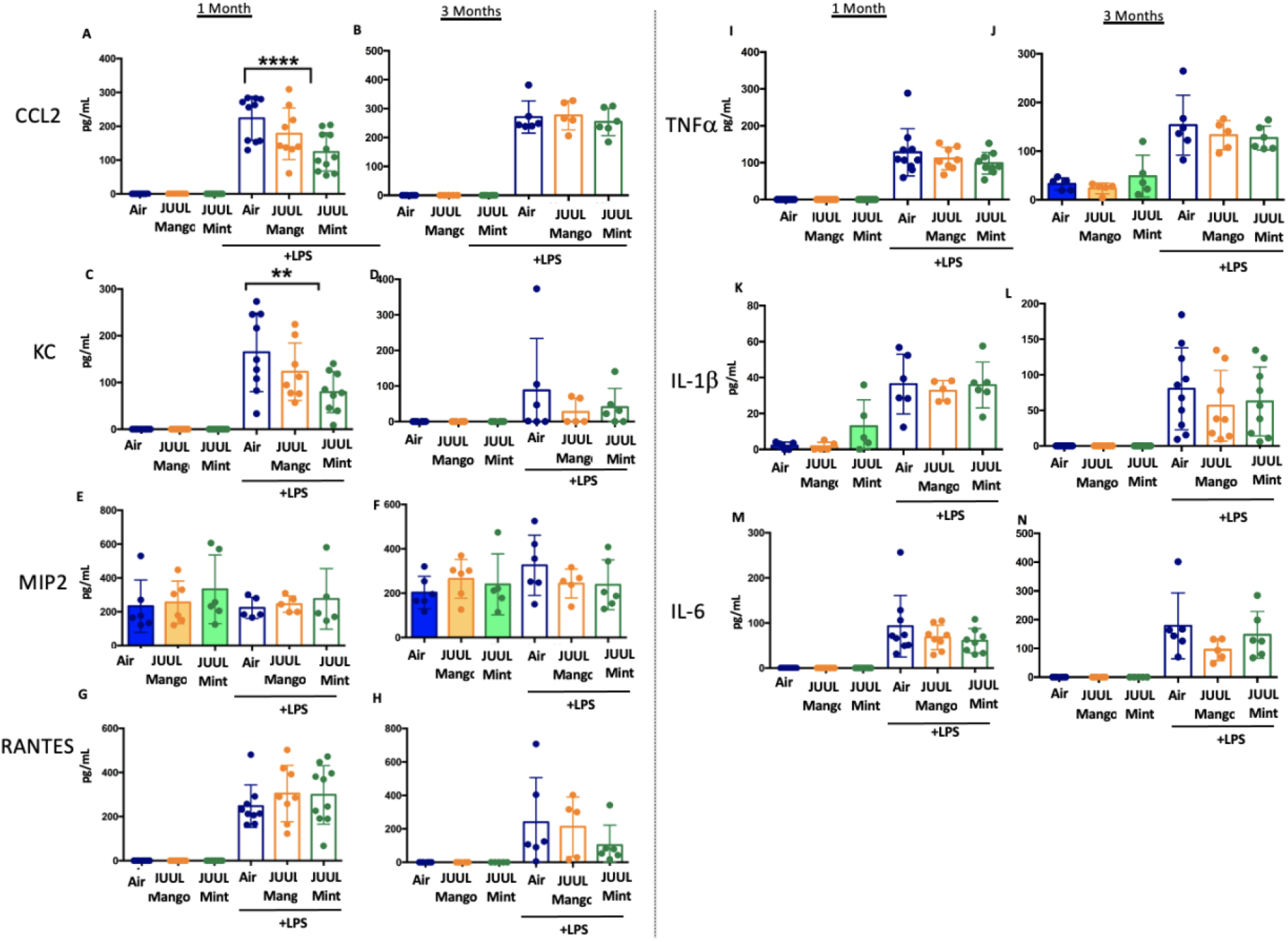
JUUL exposure alters airway inflammatory responses in the setting of inhaled LPS challenge. BAL was harvested at the endpoints, and cytokines and chemokines were quantified by ELISA. CCL2 at A) 1 month, and B) 3 months, KC at C) 1 month and D) 3 months, MIP2 at E) 1 month and F) 3 month, RANTES at G) 1 month and H) 3 months, TNFα at I) 1 month and J) 3 months, IL-1β at K) 1 month and L) 3 months, IL-6 at M) 1 month and N) 3 months. Data are presented as individual data points ± SEM with n=5-11 mice per group. \*\**p*<0.01 and **** *p*<0.0001.

### Cardiac inflammation induced by inhaled LPS challenge is increased in the setting of chronic JUUL aerosol inhalation

Bacterial pneumonia and acute lung injury lead to inflammation not only in the lungs and systemic circulation, but also in the heart (31). It is common for patients to develop myocardial inflammation and even ischemia during lung infections (31, 32). Tobacco smoking is well known to increase cardiovascular diseases and worsen outcomes in the setting of pneumonia (2, 33) and recently, it has been suggested that dual use of e-cigarettes with conventional tobacco leads to significantly higher odds of cardiovascular disease compared with cigarette smoking alone (34). Thus, we assessed the impact of acute lung injury on inflammation in cardiac tissues of JUUL exposed mice.

We assessed the expression of TNFα, IL-1β IL-6, IL-18, CCL2, CCL3, CXCL1, CXCL2, Col1a1, Col3a1, Postn and TLR4 at 1 and 3 months to determine if the LPS challenge caused changes in cardiac inflammation in the setting of JUUL aerosol inhalation (**Figure 6A-X**). LPS challenge of mice exposed to JUUL Mint for 1 month lead to significantly greater expression of cytokines (TNFα, IL-1β, IL-6) and chemokines (CCL2, CCL3, CXCL1, CXCL2) than observed in Air controls (**Figure 6A, 6C, 6E, 6I, 6K, 6M, 6O**). The enhanced chemokine induction was sustained and even further elevated after 3 month exposure to JUUL Mint (**Figure 6J, 6L, 6N, 6P**). In contrast to the elevated inflammatory response to LPS observed in mice exposed to JUUL Mint, JUUL Mango exposed mice did not have enhanced expression of cytokines or chemokines compared to Air controls. Indeed, the effects of 1 month JUUL Mint versus JUUL Mango exposure were statistically significant with regard to changes in TNFα, IL-1β, IL-18, CCL3, CXCL2 (**Figure 6A, 6B, 6G, 6K, 6M**) as well as on CXCL1 expression at both 1 and 3 months JUUL exposure (**Figure 6 M-N**).

**Figure 6.**
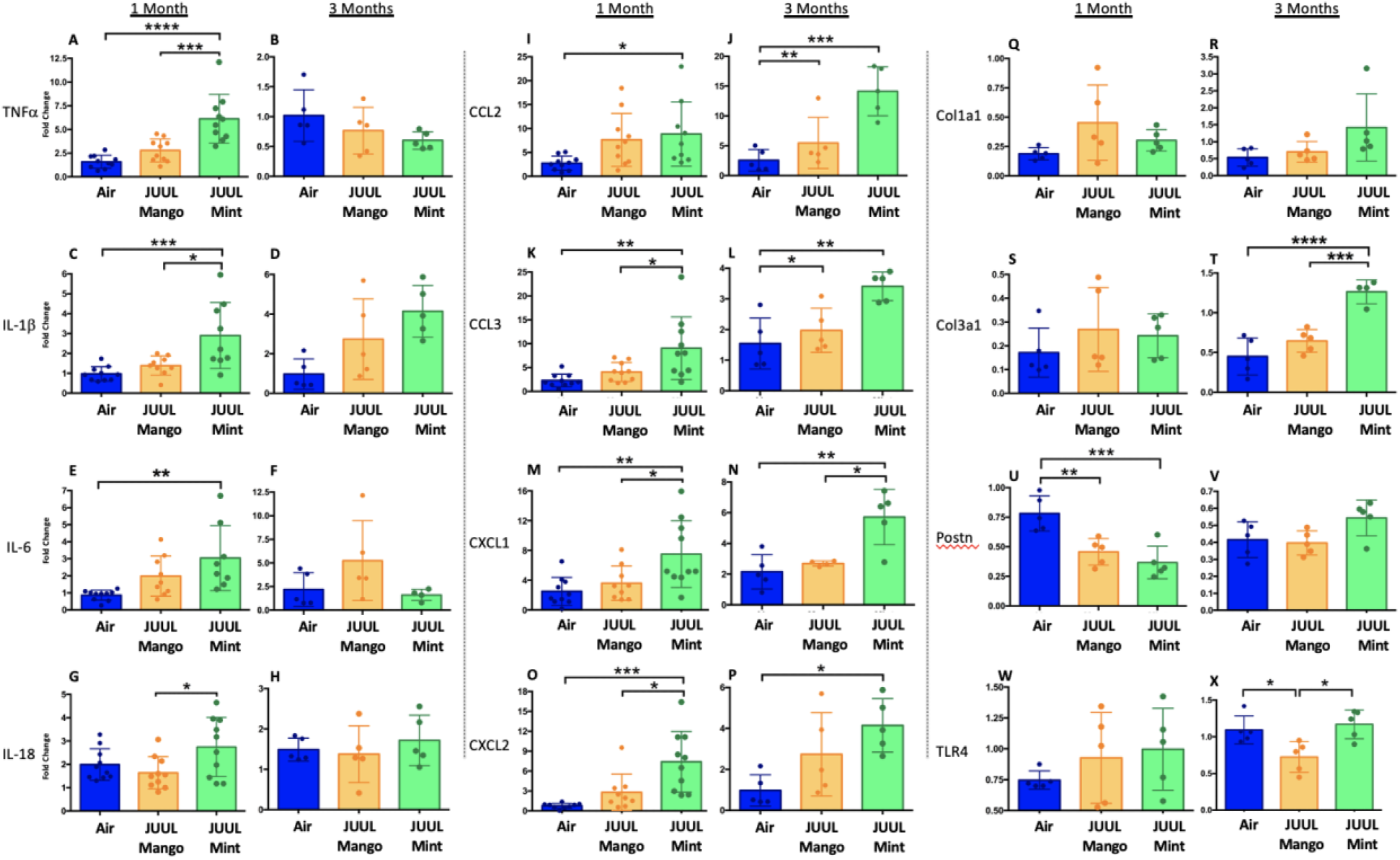
Cardiac inflammation induced by inhaled LPS challenge is increased in the setting of chronic JUUL aerosol inhalation. Hearts were harvested, and RNA was extracted from the left ventricle and qPCR was performed to quantify the gene expression of different cytokines, chemokines and fibrosis-associated genes. Cytokines include TNFα at A) 1 month and B) 3 months, IL-1β at C) 1 month and D) 3 months, IL-6 at E) 1 month and F) 3 months, and IL-18 at G) 1 month and H) 3 months. Chemokines include CCL2 at I) 1 month and J) 3 months, CCL3 at K) 1 month and L) 3 months, CXCL1 at M) 1 month and N) 3 months, and CXCL2 at O) 1 month and P) 3 months. Fibrosis-associated genes include Col1a1 at Q) 1 month and R) 3 months, Col3a1 at S) 1 month and T) 3 months, Postn at U) 1 month and V) 3 months, and TLR4 at W) 1 month and X) 3 months. Changes in expression levels are relative to Air controls. Data are presented as individual data points ± SEM with n=5-11 mice per group. \**p*<0.05, \*\**p*<0.01, \*\*\**p*<0.001 and \*\*\*\**p*<0.0001.

Enhanced inflammatory responses within tissues are known to result in fibrosis in some cases. However, analysis of pro-fibrotic gene expression only revealed that Col3a1 was significantly higher after 3 month JUUL Mint exposure (**Figure 6T**). Col1a1 and TLR4 expression were not higher in the JUUL exposed groups. Indeed, periostin expression was lower in 1 month JUUL Mint and JUUL Mango compared to Air (**Figure 6U**) and TLR4 expression was also lower in the 3 month JUUL Mango group (**Figure 6X**). Thus, while fibrotic changes are not evident, the enhanced expression of chemokines and cytokines indicates that the use of JUUL devices could predispose to cardiac tissue damage by exacerbated inflammation. In addition, we consistently found more profound effects of JUUL Mint on inflammatory cytokine and chemokine gene expression, suggesting that flavors play a significant role in heart inflammation in response to acute lung injury caused by LPS.

### Chronic JUUL inhalation by inhaled LPS challenge does not alter protein markers of the gastrointestinal tract

We showed that JUUL exposure affected the expression of pro-inflammatory cytokines in colonic tissue under homeostatic conditions. Hence, we also assessed whether this effect would be exacerbated in the context of acute lung injury. No greater increases in TNFα, IL-6 or IL-1β following LPS treatment were observed in mice subjected to 1 of 3 months of JUUL exposure (**Figure 7A-F**).

**Figure 7.**
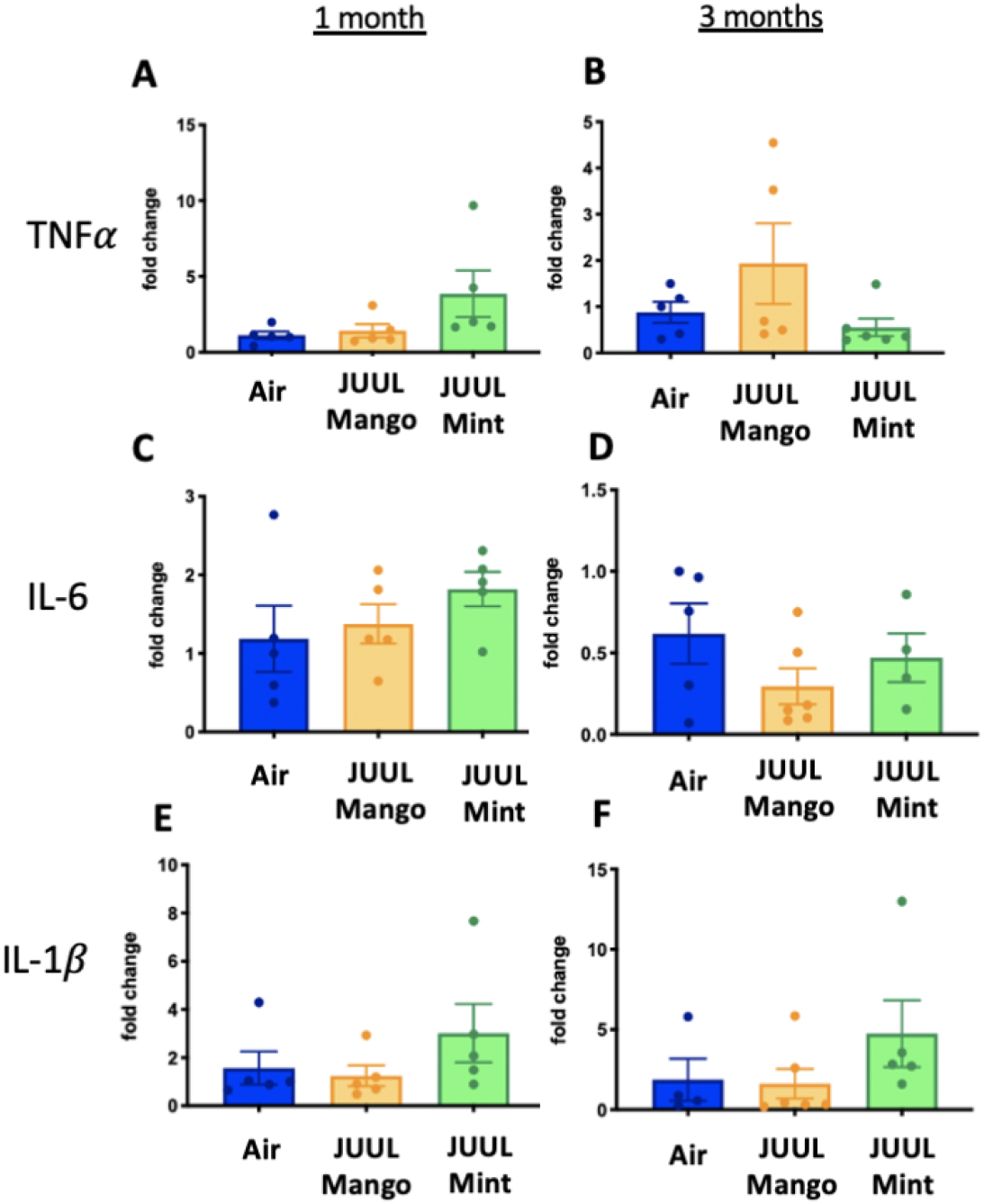
Chronic JUUL inhalation does not alter inflammatory markers in the setting of by inhaled LPS challenge in the gastrointestinal tract. Inflammation was assessed in the colon at 1 and 3 months. Panels show inflammation markers in the colon in TNF A) 1 month and B) 3 months, IL-6 at C) 1 month and D) 3 months, IL-1β at E) 1 month and F) 3 months. Data for inflammation markers are presented as individual data points ± SEM.

## Discussion

E-cigarette use has been linked to adverse cardiovascular (35) and immune responses (36, 37). However, little is known about the effects of e-cigarette use on the brain and gastrointestinal system. In this study, we found that mice exposed to flavored JUUL aerosols may induce significant neuroinflammation in the brain (**Figure 8**). The nucleus accumbens in particular was found to have elevated levels of inflammatory markers, including TNFα, IL-1β and IL-6 in both NAc-core and NAc-shell, and HMGB1-1 and RAGE in the NAc-shell (38). The NAc-core and NAc-shell contribute to the formation of anxious or depressive behaviors in the context of neuroinflammation via NFκB signaling pathway (39, 40). The NAc-core and NAc-shell are known to control reward-related behaviors through distinct neurocircuitry (41–43). We have previously shown that chronic exposure to ethanol is associated with dysregulation of the glutamatergic system and neuroinflammatory response in the NAc-shell but not in the NAc-core (44, 45). Recently, we found that 3-month exposure to either JUUL Mint or Mango induced dysregulation in glutamatergic system in the NAc-shell but not in the NAc-core (37). Our data further confirms the deleterious effects of JUUL exposure on the nucleus accumbens. On the other hand, the hippocampus, in which learning and memory are essential functions of (40), did not show significant changes in any of the cytokines tested, except for HMGB1. In the particular case of HMGB1 and its receptor RAGE, increased HMGB1 expression has been found to be a marker of neuroinflammatory conditions and may be a predictor of cognitive decline (46).

**Figure 8.**
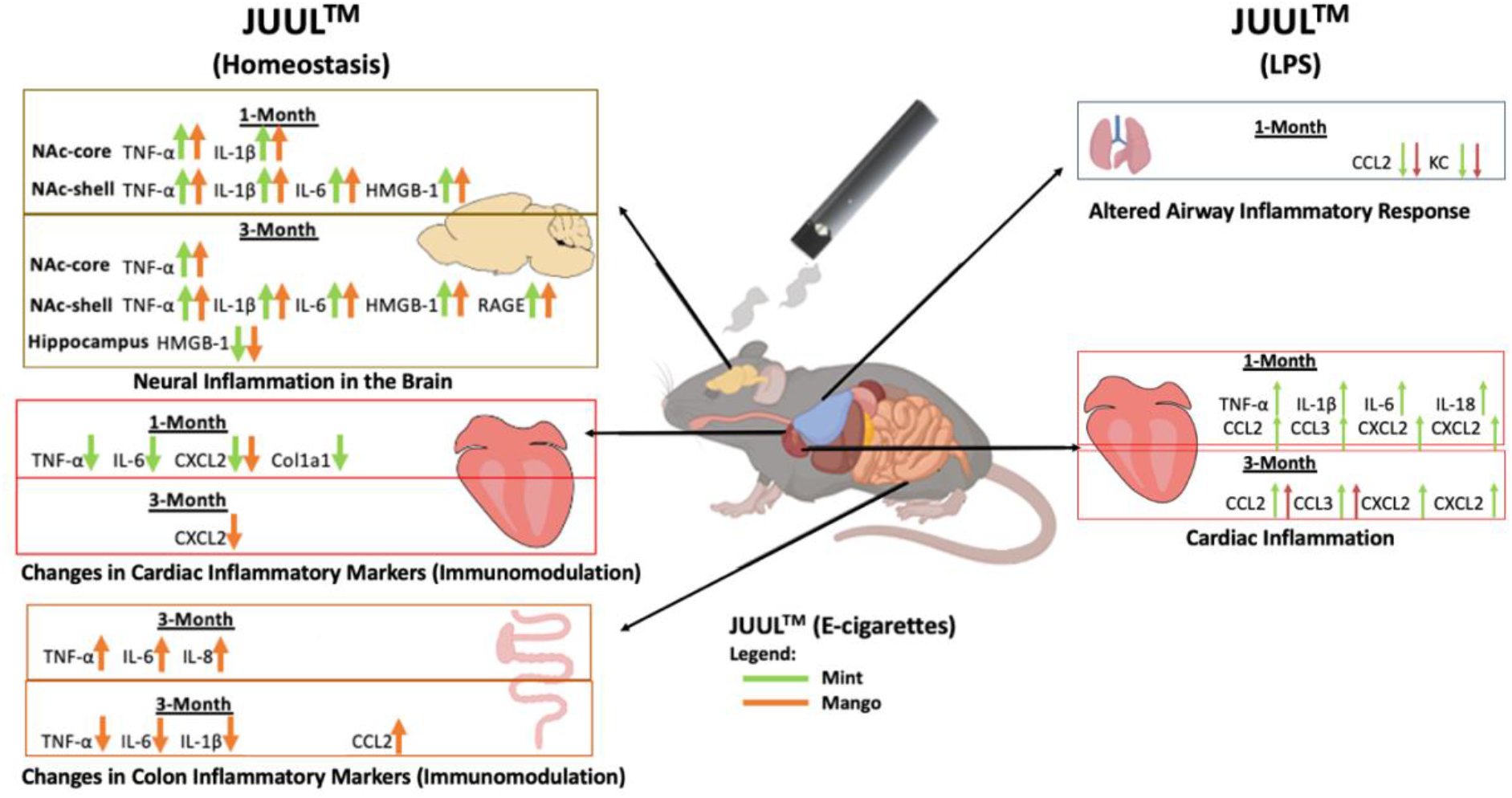
Overview of JUUL aerosol induced inflammatory changes across organs.

Exposure to drugs such as methamphetamine, cocaine and ethanol activate neuroinflammatory pathways that are associated with the release of HMGB1-1 in the striatum and nucleus accumbens, potentially linked to addictive behaviors and drug reward (45, 47). The overall similarity in inflammatory profiles and brain regions between drugs of abuse and that observed in this model of chronic JUUL exposure is certainly cause for concern as it suggests that e-cigarette use may be associated with addictive behaviors (48). Further studies into the overlap of induced neuroinflammatory pathways between drugs of abuse and JUUL is required to better understand these relationships. This includes studies in human subjects to assess the incidence of anxiety and depression in JUUL users. A correlation had previously been observed between vaping and mental health (49). Furthermore, many e-cigarette vapers, including JUUL users, are also cigarette smokers. In some cases, smokers use JUULs as an attempt to help with smoking cessation. However, data thus far demonstrates that e-cigarettes do not increase the rates of successful smoking cessation attempts (14). Indeed, neuroinflammatory effects caused by chronic, daily JUUL exposure may lead to adaptations in neural circuitry that promote addictive behaviors and drug dependence, providing a neurophysiological explanation for the observation that JUUL use does not help with smoking cessation.

It is well established that high concentrations of nicotine inhalation are toxic to the human body in a variety of ways (50). JUUL pods have been found to have the highest nicotine concentration (up to 10 times more) of any of the other cartomizer style e-cigarette or refill fluids (51). Previous studies utilizing continuous systemic delivery of nicotine via implanted pumps concluded that nicotine did not contribute to the development of neuroinflammation. The differences in findings between this study and previous work could be explained by differences in the mode of nicotine delivery, type of nicotine (free-base nicotine and nicotinic salts), inhalant device or other chemicals contained in the vaping aerosols, including vehicle components. Notably, because Mint and Mango effects differed, several of our findings point to a “non-nicotine” chemical flavorant component of the JUUL device that may be driving inflammatory changes in the brain. Recent study into the effects of vaping on the blood brain barrier lends further support to this theory, as pro-inflammatory changes were observed, partly independent of nicotine content (38).

While the nicotine concentration in JUUL pods is quite high, it does not vary with the JUUL flavor (51). The basis for the variation between the two different JUUL flavored pods tested in our study is most likely due to the differences in chemical flavorants. We observed significantly different inflammatory gene alterations in cardiac and colonic tissue in response to chronic exposure to Mint and Mango JUUL vapor (**Figure 8**). Ethyl maltol concentrations have been shown to be highest in Mango pods (1 mg/ml), while menthol concentrations are highest in cool Mint pods (10 mg/ml)(51). The most remarkable variations we observed were in response to acute lung injury through LPS challenge, where significantly higher levels of cardiac inflammatory genes were seen in mice exposed to Mint relative to Mango and controls. In the brain, inhalation of JUUL Mint aerosols led to higher TNFα and IL-1β in the NAc-shell relative to JUUL Mango. Mint aerosols are highly similar to menthol aerosols and previous studies have shown greater increases in neuronal nAChR receptors after exposure to nicotine with menthol relative to nicotine exposure alone (52). As a result, we surmise that the flavoring compound menthol in “Cool Mint” may be a factor leading to differences seen in the effects of Mint vs Mango. Overall, these findings suggest that components other than nicotine may contribute to the observed neuroinflammatory changes. Further research is needed to better understand how specific, non-nicotine JUUL components contribute to inflammatory and neuronal effects.

Collagen expression is a hallmark of fibrosis and has previously been observed in studies involving combustible cigarette smoke (30). However, compared to our prior study with Vape pens (using the nose-only InExpose system by SciReq) where profound increases in collagen deposition were observed across cardiac, hepatic and renal issues (10), we did not find increased fibrosis in these same organs in JUUL exposed mice. This raises questions about the role of different e-cigarette devices, e-liquids and experimental approaches for aerosol exposures and suggests differences in the chemical composition of aerosol and its delivery as potential causes of different biological outcomes. Research in this area is thus complex, as many teams are using a variety of devices and liquids, which may produce different effects on mammalian systems.

Intensive research has been done on the effect of cigarette smoke on inflammation and the pathogenesis of diseases such as Ulcerative Colitis and Crohn’s Disease, however there are mixed, somewhat inconclusive results when it comes to whether this exposure leads to long-term activation or suppression of inflammatory pathways, and their relation to the likelihood of developing these gastrointestinal afflictions (26). Nicotine specifically has been previously found to decrease the expression of pro-inflammatory cytokines in the colon (53). Here, we saw an increase in pro-inflammatory cytokines in the colon after 1 month of chronic exposure, whereas these same signals were significantly decreased after 3 month exposure when compared to the control group (**Figure 8**). Thus, over the course of the 3 months, the body may adapt to these changes and downregulate these markers significantly through some yet unidentified mechanism, pathway, or interaction with the specific components in the JUUL device. Whether this inflammatory adaption is beneficial or detrimental to the overall health of the colon remains to be defined.

Bacterial pneumonia and acute lung injury are known to cause inflammation systemically and in the heart (31). Indeed, effects of viral infections such as SARS-CoV-2, while originating in the lung, appear to also signal to the heart, where pathological inflammation and cardiac dysfunction are observed (54, 55). The importance of cardiac inflammation in development of heart failure following viral infection, myocardial infarction and non-ischemic cardiac injury has been increasingly appreciated (56). For example, IL-1β blockade has been shown to diminish adverse cardiac events and heart failure progression (57, 58). We demonstrate here that the hearts of mice subject to chronic JUUL exposure are significantly more sensitive to the effects of LPS delivered to the lung than are Air control mice, as evidenced by enhanced expression of pro-inflammatory cytokines and chemokines including IL-1β. The observation that there were no significant changes in vagal tone (assessed by heart rate variability) and that the pro-inflammatory enhancement by JUUL exposure was largely confined to JUUL Mint, suggests that this is not due to signals generated by direct nicotine action. While the mechanism by which chronic JUUL exposure predisposes to LPS-induced cardiac inflammation remains to be determined, these findings suggest that chronic JUUL inhalation could lead to systemic changes which sensitize maladaptive inflammatory responses that affect cardiac function.

Contrary to our initial expectations, we did not find significant changes in autonomic tone or pulmonary function with daily, long-term JUUL aerosol exposure. Our model is limited in that mice are primarily nose breathers and we used whole-body exposure, so it is possible that the extent of e-cigarette aerosol exposure at the level of the alveoli may be lower than in humans due to aerosol deposition within the nasal cavity. Alternatively, our study may be underpowered to detect subtle differences induced by JUUL vaping. However, it is important to mention that this is the first study assessing JUUL devices in a multiorgan fashion. In addition, we found clear effects of JUUL aerosols on inflammatory responses in organs other than the lungs. Thus, the effects of e-cigarette exposure may be greater on organs far removed from the lungs, the first organ to come in contact with aerosols.

## Conclusion

These data indicate that chronic inhalation of chemicals within e-cigarette aerosols leads to identifiable inflammatory changes within multiple organs. JUUL users may unwittingly expose themselves to neurologic, colonic and cardiac inflammatory effects. Further research is needed to better understand the long-lasting effects of vaping.

## Materials and Methods

### JUUL Exposures

Six to eight week old female C57BL/6 mice were purchased from Envigo. Mice were placed in individual sections of a full-body exposure chamber (Scireq) for 20 minutes daily for 5 days per week, for 4-12 weeks. Mice were exposed to either e-cigarette aerosol created from Mango JUUL pods or Mint JUUL pods containing 5% nicotinic salts (59 mg/ml) using the InExpose system (Scireq). Air control mice were placed in an identical chamber for the same amount of time but inhaled room air only. A 3-D printed adapter was created to produce a tight fit for the JUUL device (designed and produced by Vitorino Scientific LLC). A negative pressure of 2L/s was used to activate the e-cigarette for 4 seconds followed by 16 seconds of room air at 2L/s. The final exposure was done 30 minutes prior to harvest. All experiments were conducted with approval of the UCSD Institutional Animal Care and Use Committee (IACUC protocol S16021). All authors complied with the ARRIVE guidelines.

### LPS Intranasal Challenge

Mice were sedated with isoflurane, held upright and intranasally challenged with LPS *(E.coli* O111:B4; Sigma) at a concentration of 2.5 μg per gram of mouse in 0.9% saline (100 μl). The LPS challenge was given through the left nare to decrease liquid trapping in the nasopharyngeal dead space. Mice were maintained in the upright position until respirations returned to normal. Mice were monitored overnight prior to harvest 24 hours after challenge.

### Assessment of Pulmonary Function

At the end of 4 and 12 weeks of exposure, prior to harvest, mice were sedated via intraperitoneal (i.p.) injection of ketamine 10mg/ml xylazine 100mg/ml. Mice underwent tracheostomy with 18g metal cannula and were attached to the FlexiVent mouse ventilator (SciReq). Measurements of lung physiology via mechanic scans were obtained at baseline, followed by assessment of physiologic responses to methacholine (MCH) challenge at 0, 6, 12 and 24 mg/ml, including Respiratory Resistance (Rrs) and Elastance (Ers). Pressure Volume (PV) loops were also obtained.

### Cell Counts and Differential

Bronchoalveolar lavage (BAL) was collected by flushing airways with ice cold 800 ul PBS three times via mouse tracheal cannulation. Samples were pelleted at 3000 rpm for 4 minutes at 4°C. Pellets were resuspended in 1 ml of ice cold PBS, counted with Countess (Life Technologies) for total cells quantification. Later, two dilutions (1:1 and 1:4) of total cells were cytospun onto slides at 800 rpm for 3 minutes and then cells were fixed with Giemsa Wright. Slides were de-identified and randomized prior to blinded cell counting; 200 cells from each slide were counted via light microscopy under 40X magnification, and finally percentage of neutrophils was calculated and total amounts of neutrophils extrapolated based on total cell quantification.

### Histology

Lungs were inflated with Zfix (Anatech ltd) at 25cm H2O pressure for 18 hours, followed by transfer into 75% ethanol prior to paraffin embedding. Lung slices were stained with H&E.

### Fibrosis Analysis

The right kidney, one lobe of liver, and the base of the heart were then immediately dissected after euthanasia and placed in Z-fix at 4°C. After 48 h, all organs were moved to 75% ethanol and submitted to the University of California, San Diego histology core for paraffin embedding. Collagen was detected in 5-μm sections first by Masson’s trichrome stain. All histology slides underwent quantification of fibrosis by calculating the mean percent fibrotic area in 15-25 randomly acquired ~20 images using computer-aided morphometry performed using ImageJ. Briefly, using the color threshold with default thresholding method, red threshold color and HSB color space, the total area of tissue in the slide was selected and measured, later the tissue stained for Masson’s trichrome blue was also selected and measured prior adjustment of the “Hue” parameter (Saturation Brightness/Value Each colour shade). Then, a percentage of the area stained by Masson’s trichrome blue was determined relative to the total tissue area. All histology slides from the same tissue group were blinded and underwent these computer analyses in an identical fashion. Fibrotic area is presented relative to that of air controls.

### Isolation of RNA from the murine colonic tissue and qRT-PCR for inflammatory cytokines

RNA was isolated from mouse colon tissues using the Zymo miniprep kit according to the manufacturer’s instructions, followed by cDNA synthesis. Quantitative Real-Time PCR was conducted for target genes and normalized to housekeeping gene 18S rRNA. Primer sequences are provided in **Table 1**.

**Table 1.**
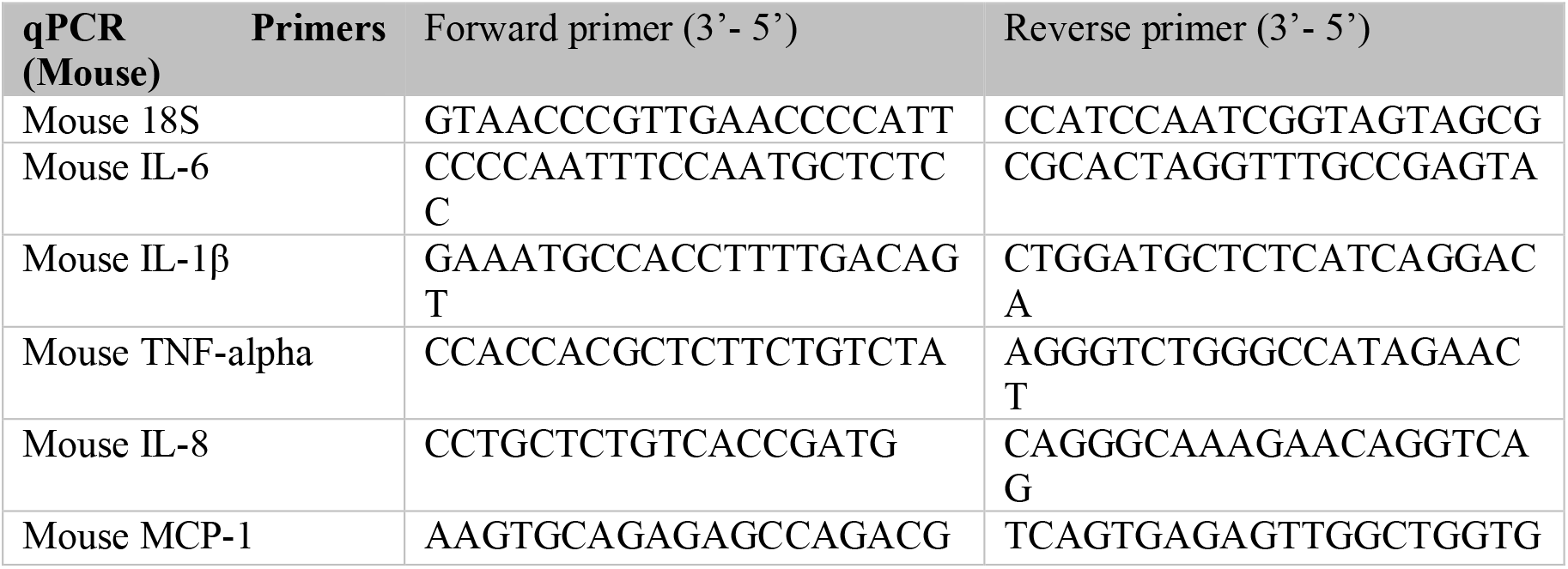

### Cardiovascular Physiology Measurements

Heart rate, heart rate variability (HRV) and blood pressure measurements were taken after the last exposure to JUUL aerosol or Air at 1 and 3 months, via the Emka non-invasive ECG Tunnels and the CODA non-invasive blood pressure system. Prior to data collection, mice were acclimated for 20 minutes per day for the last 3 days in the ECG and blood pressure systems. Heart rate variability was determined through time-domain measurements, specifically SDNN and RMSSD. The SDNN is the standard deviation of all normal R-R intervals, providing information on total autonomic variability. The RMSSD is the root mean square of those standard deviations and represents the variability in the short term.

### Brain tissue harvesting

At the end of the experiments, mice were euthanized by ketamine and xylazine i.p. injection, rapidly decapitated, with their brains removed and stored at −80°C. The cryostat apparatus maintained at −20°C and used to dissect NAc-core, NAc-shell, and HIP, which micropunched stereotaxically. The stereotaxic coordinates for the mice brain (59) was used to isolate the brain regions of interest following visualized landmarks.

### qRT-PCR

Total RNA from the NAc-core, NAc-shell and HIP of JUUL Mango, JUUL Mint exposed groups, in addition to Air control group. Brain tissue was extracted with TRIzol reagent, using the manufacturer’s protocol (Invitrogen, USA). The cDNA was synthesized using the iScript cDNA synthesis kit (Bio-Rad, USA). The mRNA expression level of the brain tissue was detected by qRT-PCR via iQ SYBER green I Supermix (Bio-Rad, USA) and a Bio-Rad RT-PCR instrument system. The thermocycling protocol consisted of 10 min at 95°C, 40 cycles of 15 sec at 95°C, 30 sec at 60°C, and 20 sec at 72°C and completed with a melting curve ranging from 60–95°C to facilitate distinction of specific products. A reaction with primers of TNFα, IL-1β and IL-6 was performed, the glyceraldehyde-3-phosphate dehydrogenase (GAPDH) gene was used as a housekeeping control. Data were expressed as fold change (2^-δδ^Ct) relative to the control group. The primer sequences are listed in **Table 2**.

**Table 2.**
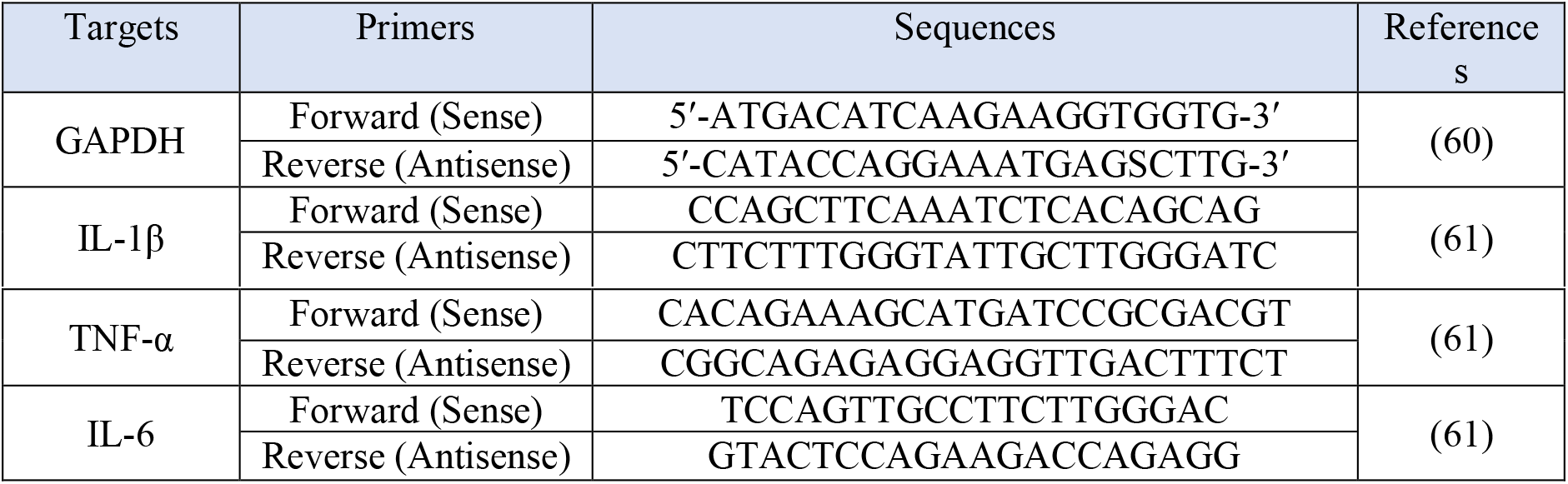

### Brain Western Blot

Immunoblot assays were conducted to measure the expression of HMGB1 and RAGE proteins in the NAc core, NAc shell and HIP as described previously (62). Briefly, the samples were homogenized with lysis buffer containing protease and phosphatase inhibitors. The amount of protein in each tissue sample was quantified using detergent compatible protein assay (Bio-Rad, Hercules, CA, USA). Then, 10% polyacrylamide gels used, in which, an equal amount of protein from each sample was loaded. Proteins were then transferred to a PVDF membrane and blocked with 5% fat free milk in Tris-buffered saline with Tween-20 (TBST). Membranes then incubated with appropriate primary antibodies at 4°C (overnight): rabbit anti-HMGB1 (1:1000; Abcam), rabbit anti-RAGE (1:1000; Abcam) and mouse anti-β-tubulin (1:1000; BioLegend; used as a control loading protein). Membranes were then incubated with appropriate secondary antibody (1:5000) for 90 minutes at room temperature. Chemiluminescent reagents (Super Signal West Pico, Pierce Inc.) were incubated with the membranes. The GeneSys imaging system was used and the digitized blot images were developed. Quantification and analysis of the expression of HMGB1, RAGE and β-tubulin were performed using ImageJ software. Air control group data were represented as 100% to assess the change in the expression of the protein of interest as described previously (63).

### RNA Isolation, cDNA Synthesis and qRT-PCR from Cardiac Tissue

RNA was isolated from samples of cells or tissue homogenized in TRIzol (Invitrogen), with subsequent extraction with chloroform, precipitation of RNA with isopropanol, and washing of RNA pellet twice with 70% ethanol. Synthesis of cDNA from isolated RNA was carried out using the High-Capacity cDNA Reverse Transcription kit with RNase inhibitor (Applied Biosystems). qRT-PCR was carried out using predesigned PrimeTime qPCR Primers (IDT) and TaqMan Universal Master Mix II with UNG (Applied Biosystems), combined with cDNA samples in a 96-well PCR plate and run on a 7500 Fast Real–Time PCR system (Applied Biosystems). The gene expression data acquired was analyzed using the comparative 2^-ΔΔCT^ method, with GAPDH expression levels used as the internal control.

### Cytokine Profiling

Cytokine and chemokine levels were assessed in the BAL with Duo-Set Enzyme-Linked Immunosorbent Assays (R&D Systems Inc., Minneapolis, MN). ELISAs were performed per manufacture’s instructions.

### Statistical Analyses

Analyses were conducted using GraphPad Prism v6.0 or v8.0. Assays with data from more than 2 groups or timepoints were analyzed by ANOVA and are presented as means +/− SEM. Quantification and analysis of Western blot protein levels of HMGB1, RAGE and β-tubulin, and histologic examination of tissue fibrosis, were performed using ImageJ software. The gene expression data acquired by qPCR was analyzed using the comparative 2^-ΔΔCT^ method, with GAPDH (brain) and 18S rRNA (colon) expression levels used as the internal control.

## Author contributions

Conception and design of the experiments: AM, CB, HA, JS, JAMS, AS, DA, SD, ZS, YS, JHB and LCA. Acquisition of data: AM, CB, HA, JS, IA, DG, AS, SM, AJ, SN, JP, SP, KP and RAK. Analysis and interpretation of data: AM, CB, HA, JAMS, AS, DA, SD, MB, ZS, YS, JHB and LCA. Manuscript composition: AM, CB, HA, JAMS, IA, DG, SD, YS, JHB and LCA. All authors reviewed, contributed to, and approved the manuscript.

## Data and materials availability

Please contact Dr. Crotty Alexander.

