## Supplemental Data for "Effect of chronic JUUL aerosol inhalation on inflammatory states of the brain, lung, heart and colon in mice"

### Supplemental Figures

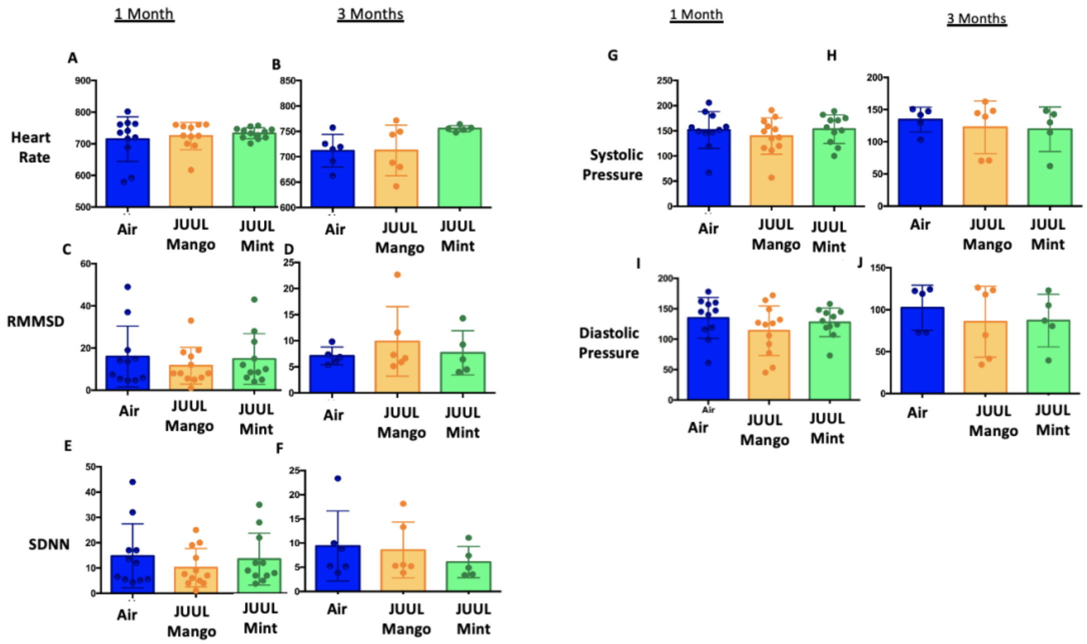


**Supplemental Figure 1. JUUL aerosol inhalation does not alter heart rate, heart rate variability or blood pressure.** Before assessment of lung function, mice underwent heart rate and blood pressure measurements using Emka non-invasive ECG Tunnels and the CODA non-invasive blood pressure system at 1 and 3 months. Heart rate (A-B), heart rate variability as measured by RMMSD and SDNN (C-F), systolic blood pressure (G-H) and diastolic blood pressure (I-J) were unaltered by chronic e-cigarette aerosol inhalation. Data are presented as individual data points ± SEM with n=5-11 mice per group.

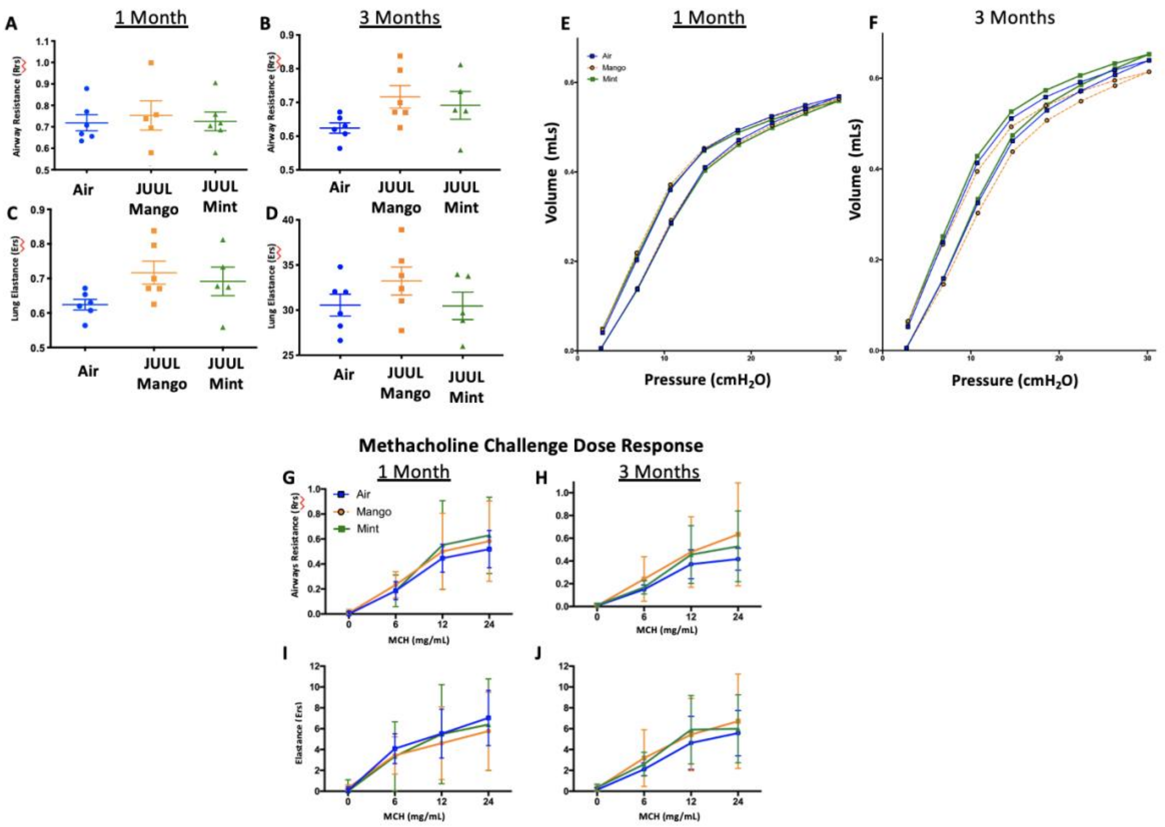


**Supplemental Figure 2. Chronic exposure of JUUL does not increase airways resistance or induce airways hyperreactivity.** At end points prior to harvest, mice underwent tracheostomy and attached to the FlexiVent mouse ventilator (SciReq). Airways resistance, lung elastance and pressure-volume (PV) loops were measured by mechanics scans using the FlexiVent mouse ventilator (SciReq)(A-F), followed by the assessment of responses to methacholine (MCH) challenge at 0, 6, 12 and 24 mg/mL (G-J). These parameters were assessed after 1 and 3 months of exposure of JUUL. Panels show airways resistance (Rrs) at A) 1 month and B) 3 months, Elastance at C) 1 month and D) 3 months, and PV loops at E) 1 month and F) 3 months. There were no differences in responses to methacholine challenge by Rrs at G)1 month and H) 3 months, or Ers at I) 1 month and J) 3 months. Data for airways resistance and lung elastance are presented as individual data points ± SEM with n=5-11 mice per group, and PV loops as means with n=6 mice per group.

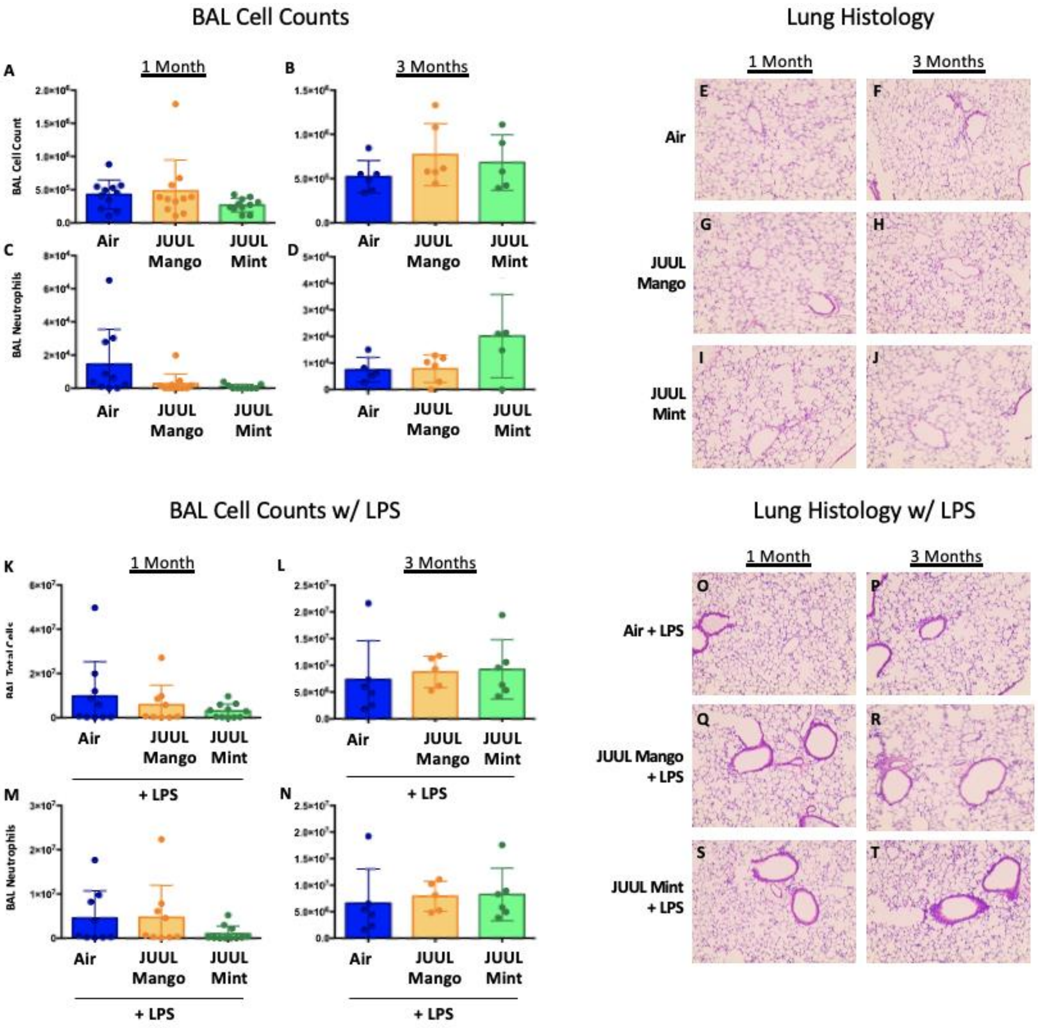


**Supplemental Figure 3. Chronic JUUL exposure does not affect leukocyte levels in the airways and lung parenchyma at baseline or influx into the airways and parenchyma in the setting of inhaled LPS challenge.** BAL was obtained, leukocyte counts were performed and the left lung lobe was fixed with formalin at 25 $cm^3$ water pressure and stained with H&E. BAL total cell counts in air versus JUUL exposed mice were no different at A) 1 month and B) 3 months, nor were neutrophils counts at C) 1 month and D) 3 months. Representative pictures from H&E staining of lung tissue are shown in E,G,I) for 1 month and F,H,J) 3 months. In the setting of acute lung injury induced by inhaled LPS challenge, cell counts in the BAL were no different across groups (K-N), nor was lung inflammation by histologic evaluation (O-T). Data for cell counts are presented as individual data points $\pm$ SEM and representative pictures for H&E lung stained tissue with n=5-11 mice per group.

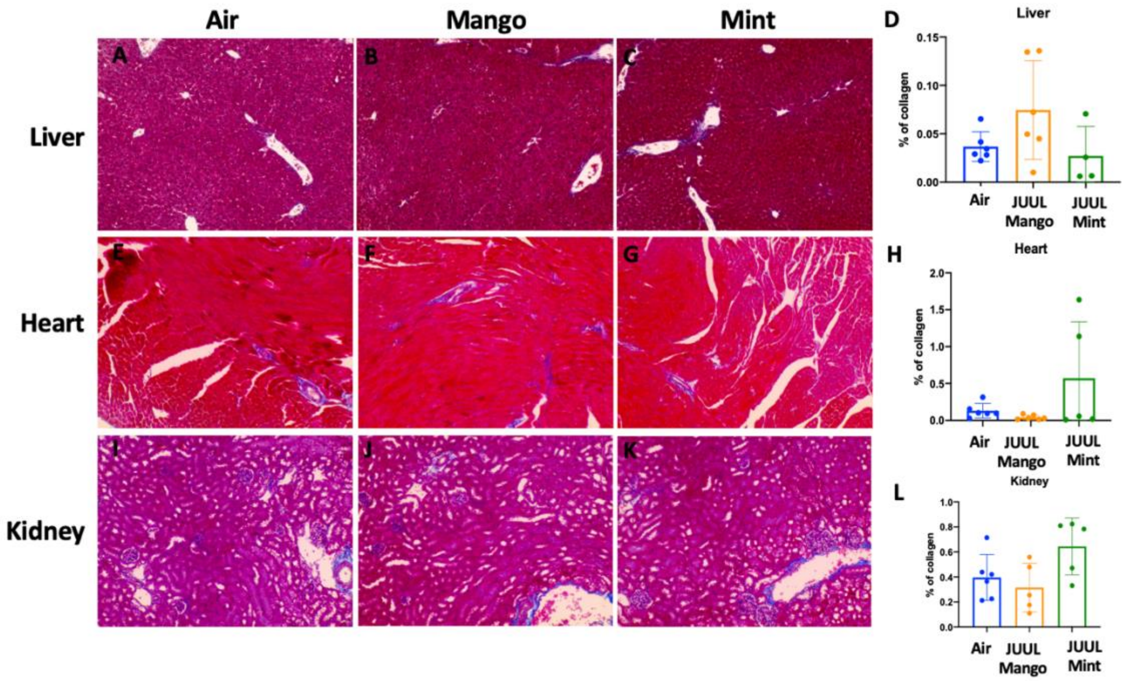


**Supplemental Figure 4. Chronic exposure of JUUL for 3 months does not induce fibrosis in the liver, heart and kidney.** Collagen deposition was quantified by image analysis (using ImageJ) of lung histological slides stained with Masson's trichrome. Representative pictures are shown for liver tissue in A) Air control, B) JUUL Mango, C) JUUL Mint and D) Quantification of collagen percentage in liver tissue. Representative pictures for heart tissue are shown in E) Air control, F) JUUL Mango, G) JUUL Mint and H) Quantification of collagen percentage in heart tissue. Representative pictures for heart tissue are shown in I) Air control, J) JUUL Mango, K) JUUL Mint and L) Quantification of collagen percentage in kidney tissue. Data for quantification is presented as individual data points ± SEM with n=4-6 mice per group.
